# TCF7L2: a potential key regulator of antidepressant effects on hippocampal astrocytes in depression model mice

**DOI:** 10.1101/2023.08.09.552412

**Authors:** Yusaku Koga, Naoto Kajitani, Kotaro Miyako, Hitoshi Takizawa, Shuken Boku, Minoru Takebayashi

## Abstract

**Aim:** Clinical and preclinical studies suggest that hippocampal astrocyte dysfunction is involved in the pathophysiology of depression; however, the underlying molecular mechanisms remain unclear. Here, we attempted to identify the hippocampal astrocytic transcripts associated with antidepressant effects in a mouse model of depression.

**Methods:** We used a chronic corticosterone-induced mouse model of depression to assess the behavioral effects of amitriptyline, a tricyclic antidepressant. Hippocampal astrocytes were isolated using fluorescence-activated cell sorting, and RNA sequencing was performed to evaluate the transcriptional profiles associated with depressive effects and antidepressant responses.

**Results:** Depression model mice exhibited typical depression-like behaviors that improved after amitriptyline treatment; the depression group mice also had significantly reduced GFAP-positive astrocyte numbers in hippocampal subfields. Comprehensive transcriptome analysis of hippocampal astrocytes showed opposing responses to amitriptyline in depression group and control mice, suggesting the importance of using the depression model. Transcription factor 7 like 2 (TCF7L2) was the only upstream regulator gene altered in depression model mice and restored in amitriptyline-treated depression model mice. In fact, TCF7L2 expression was significantly decreased in the depression group. The level of TCF7L2 long non-coding RNA, which controls mRNA expression of the TCF7L2 gene, was also significantly decreased in this group and recovered after amitriptyline treatment. The Gene Ontology biological processes associated with astrocytic TCF7L2 included proliferation, differentiation, and cytokine production.

**Conclusion:** We identified TCF7L2 as a gene associated with depression- and antidepressant-like behaviors in response to amitriptyline in hippocampal astrocytes. Our findings could provide valuable insights into the mechanism of astrocyte-mediated antidepressant effects.

## Introduction

According to the “Global Health Estimates” released recently by the World Health Organization, approximately 4.4% of the world’s population suffers from major depressive disorder (MDD), which is ranked as the largest contributor to non-fatal health loss. The monoamine hypothesis is used most commonly to explain the pathophysiology of MDD, and most antidepressants have been developed based on this hypothesis; however, it cannot explain why up to 30% of patients with MDD are resistant to currently available antidepressants. Additionally, it is increasingly becoming clear that non-neural glial cells play an important role in the regulation of various neural activities.

Astrocytes are the most abundant and versatile of all glial cell types and are crucial to the neuronal microenvironment due to their involvement in regulating glucose metabolism, uptake of neurotransmitters such as glutamate, synaptic development and maturation, and blood-brain barrier (BBB) functioning.^1^ We have previously reported that amitriptyline (AMI), a tricyclic antidepressant (TCA), acts directly on astrocytes and increases the levels of various neurotrophic factors associated with antidepressant effects, suggesting that at least some of these effects are astrocyte-mediated^2, 3^ and that astrocytes may play important roles in the pathophysiology of depression. A growing number of reports suggest that hippocampal astrocytes in particular are involved in the pathogenesis and treatment of depression.^4–9^ Hippocampal astrocytes are closely related to the hypothalamic-pituitary-adrenal (HPA) axis, and elevated glucocorticoid levels associated with the disruption of the HPA system can reduce astrocyte numbers.^10, 11^ Postmortem studies have shown consistent reductions in the density of astrocytes immunoreactive for glial fibrillary acidic protein (GFAP) in patients with MDD.^1^ Furthermore, in postmortem brain studies focusing on the hippocampus, the density of GFAP-immunoreactive astrocytes was found to be decreased in depression only in the absence of antidepressants.^12^ It has also been reported that astrocytes in the dentate gyrus and cornu ammonis 3 (CA3) regions of the hippocampus respond to antidepressants, as the administration of fluorocitrate (a reversible inhibitor of astrocyte function) to these regions was found to inhibit the antidepressant effect of imipramine, another TCA, in a mouse model of depression.^13^ Moreover, antidepressants affect not only a variety of astrocyte-related proteins and mRNA expression in the hippocampus, but also influence gliogenesis.^5^

Despite hippocampal astrocytes being implicated in the pathogenesis of MDD and in the effects of antidepressants, the underlying molecular mechanisms have not been elucidated. An important reason for this could be that many studies have focused on analyzing brain tissue samples with various cell types, which may have masked the specific contributions of astrocytes. Furthermore, there have been very few integrated studies on astrocytes with regard to both depression onset and antidepressant effects. Thus, the present study was designed to explore molecules that correlate with behavior in a more clinically relevant mouse model of depression, wherein depression-like behavioral abnormalities are appropriately corrected by antidepressant treatment.

## Methods

### Experimental animals

Wild-type male C57BL/6J mice were purchased from Jackson Laboratory Japan, Inc. (Kanagawa, Japan). All mice were housed under a 12-h light/dark cycle (lights on, 8 am– 8 pm) at a controlled temperature (23–25 °C) and with ad libitum access to food and water. Mice were allowed to habituate without experimentation for one week after being brought in. Handling was performed at least once a week for at least 5 min until the end of the study. All procedures involving experimental animals were approved by the Institutional Animal Committee of Kumamoto University (Recognition No. A2022-045R1) and performed in accordance with their guidelines.

The mouse model was established as previously described^14^. Briefly, a 35 μg/mL aqueous solution of corticosterone (CORT; C0388; TCI, Tokyo, Japan) in 0.45% w/v 2-hydroxypropyl-β-cyclodextrin (324-84233; FUJIFILM, Tokyo, Japan) as a solubilizing agent and vehicle was prepared. As shown in Fig. 1a, 8-week old mice were randomly assigned to either the corticosterone treatmented group (DEP) or vehicle treatmented group (CON). After 1 week of acclimation, CORT solution or vehicle was administered to the mice daily for 7 weeks. From week 4, the mice were randomly divided into two groups, one receiving AMI (10 mg/kg; A0908; TCI) and the other receiving saline as the control by intraperitoneal injection. The AMI treatment groups were defined as AMI-treated CON (CON+AMI) and AMI-treated DEP (DEP+AMI). Doses were administered 6 days per week (on all days except for the day of the sucrose preference test [SPT]).

**Figure 1.**
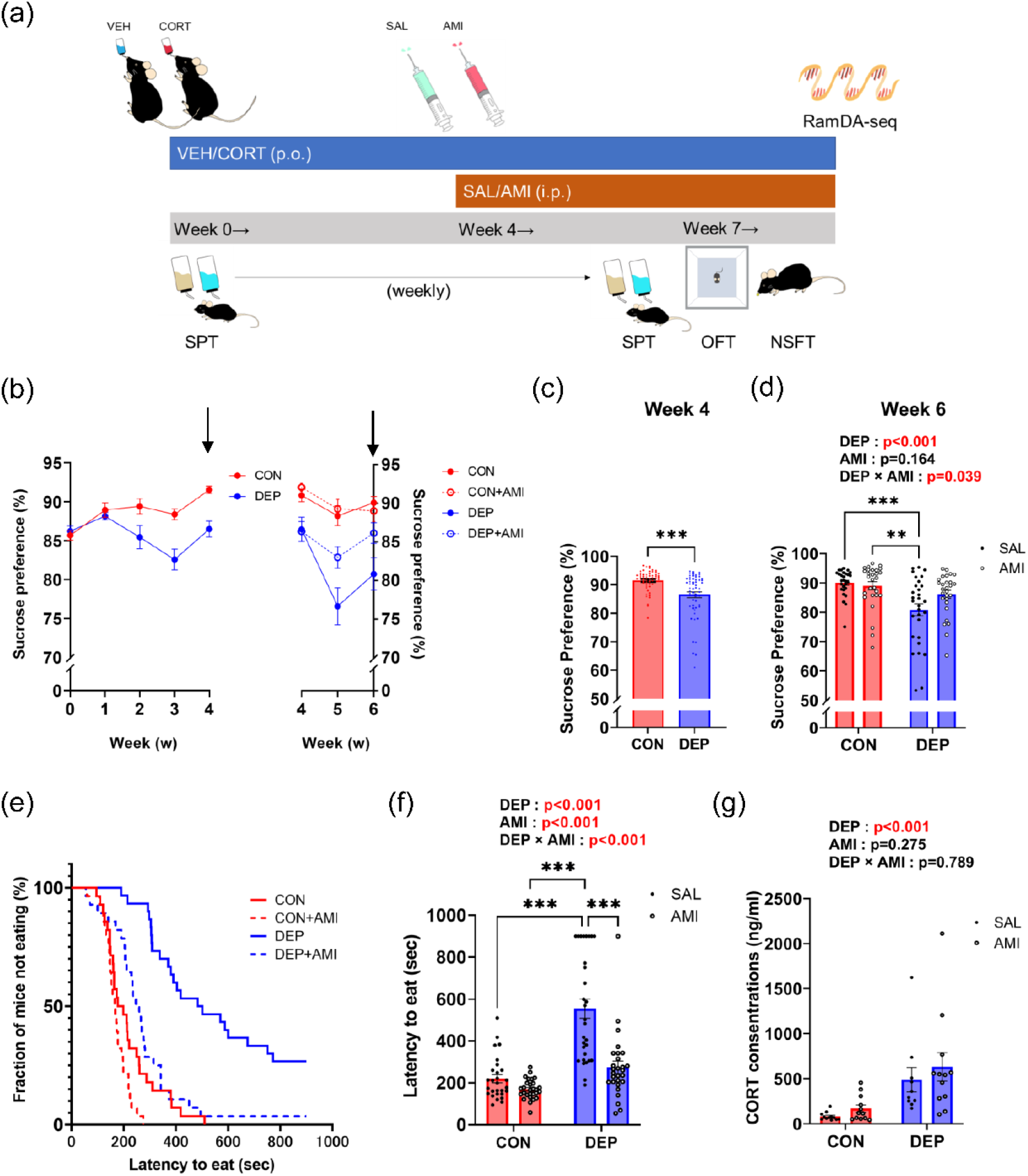
Establishment of a mouse model of CORT-induced depression for assessing antidepressant effects. (a) A schematic outlining the study design and experimental manipulations. CORT or vehicle was administered daily to acclimated mice over 7 weeks, with saline or AMI intervention initiated from week 4. n = 28–30 mice/group. (b) Preference values in the SPT over time. Results are shown in Figs. 1c and 1d just before AMI administration started (week 4) and two weeks after administration started (week 6). (c) The SPT preference value in week 4 was significantly lower in the DEP group (t (112) = 4.418, *p* < 0.001, unpaired t-test). (d) Sucrose preference was improved to the basal level in the DEP+AMI group (DEP ☓AMI: F (1, 110) = 4.360, *p* = 0.039, two-way ANOVA). (e) NSFT results are shown in terms of a Kaplan-Meier cumulative survival curve. The latency to feed was prolonged in the DEP group, and showed improvement to the base level in the DEP+AMI group. (f) Scatter graph depicting individual latency to feed. The test was finished at 900 seconds, and for mice that exceeded that time, the time to feed was analyzed as 900 seconds. The DEP group had a significantly prolonged latency to feed (F (1, 110) = 52.68, *p* < 0.001, two-way ANOVA), which was shortened in the DEP+AMI group (DEP☓AMI: F (1, 110) = 14.45, *p* < 0.001, two-way ANOVA). (g) Corticosterone concentration was elevated in the DEP group and was not affected by AMI treatment (DEP: F (1, 42) = 17.26, *p* < 0.001; AMI: F (1, 42) = 1.222, *p* = 0.275; DEP☓AMI: F (1, 42) = 0.073, *p* = 0.789, two-way ANOVA). n = 10–12 mice/group. CORT: corticosterone, AMI: amitriptyline CON: control, DEP: depression model mice, CON+AMI: AMI-treated CON, DEP+AMI: AMI-treated DEP. **, *p* < 0.01; ***, *p* < 0.001.

### Behavioral tests

The SPT and novelty suppressed feeding test (NSFT) were conducted to check for the appearance or improvement of depression-like behavior. The SPT is used to assess anhedonia, a core symptom of depression in humans, defined as the inability to experience pleasure from rewarding or enjoyable activities.^15^ The NSFT is used to assess anxiety-induced hyponeophagia—reduced feeding upon exposure to a novel environment is another core symptom of depression, and chronic administration of antidepressants is known to reduce the time to food intake.^16^ The open field test (OFT) was conducted to assess locomotor activity^17^ to confirm that CORT and AMI were not exerting a strong influence on the animals’ activity levels. More information regarding the behavioral experiments is provided as Supporting Information.

### Enzyme-linked immunosorbent assay (ELISA)

To minimize fluctuations in CORT concentration, mice were bled from 09:00 a.m. to 11:00 a.m. Blood samples were collected with ethylenediaminetetraacetic acid (EDTA) as an anticoagulant, and EDTA-plasma was isolated by centrifugation (1,500 g, 4 °C, 15 min) before storage at −30 °C until further use. Plasma CORT concentrations were measured using the Corticosterone AssayMax ELISA Kit (EC3001-1; Assaypro, St. Charles, USA) following the manufacturer’s instructions. A 4-parameter logistic curve was prepared using standard products, and sample optical densities at 450 and 570 nm were quantified using a microplate spectrophotometer (Varioskan LUX; Thermo Fisher Scientific, Waltham, MA, USA). The OD value for each diluted sample was calculated by subtracting the measured value at 570 nm from that at 450 nm to correct for optical defects.

### Immunohistochemistry and imaging

Mice were perfusion-fixed using 4% paraformaldehyde and then sacrificed. Frozen sections were prepared, mounted on glass slides, and immunostained using standard methods; more details are provided as Supporting Information. Images were captured using a fluorescence microscope (BZ-X700; Keyence, Osaka, Japan). Images of the bilateral ventral hippocampus were captured from seven to eight mice per group at 40× magnification. The number of GFAP-positive cells and the GFAP-positive area per cell in each subfield (Fig. 2a) were measured.

**Figure 2.**
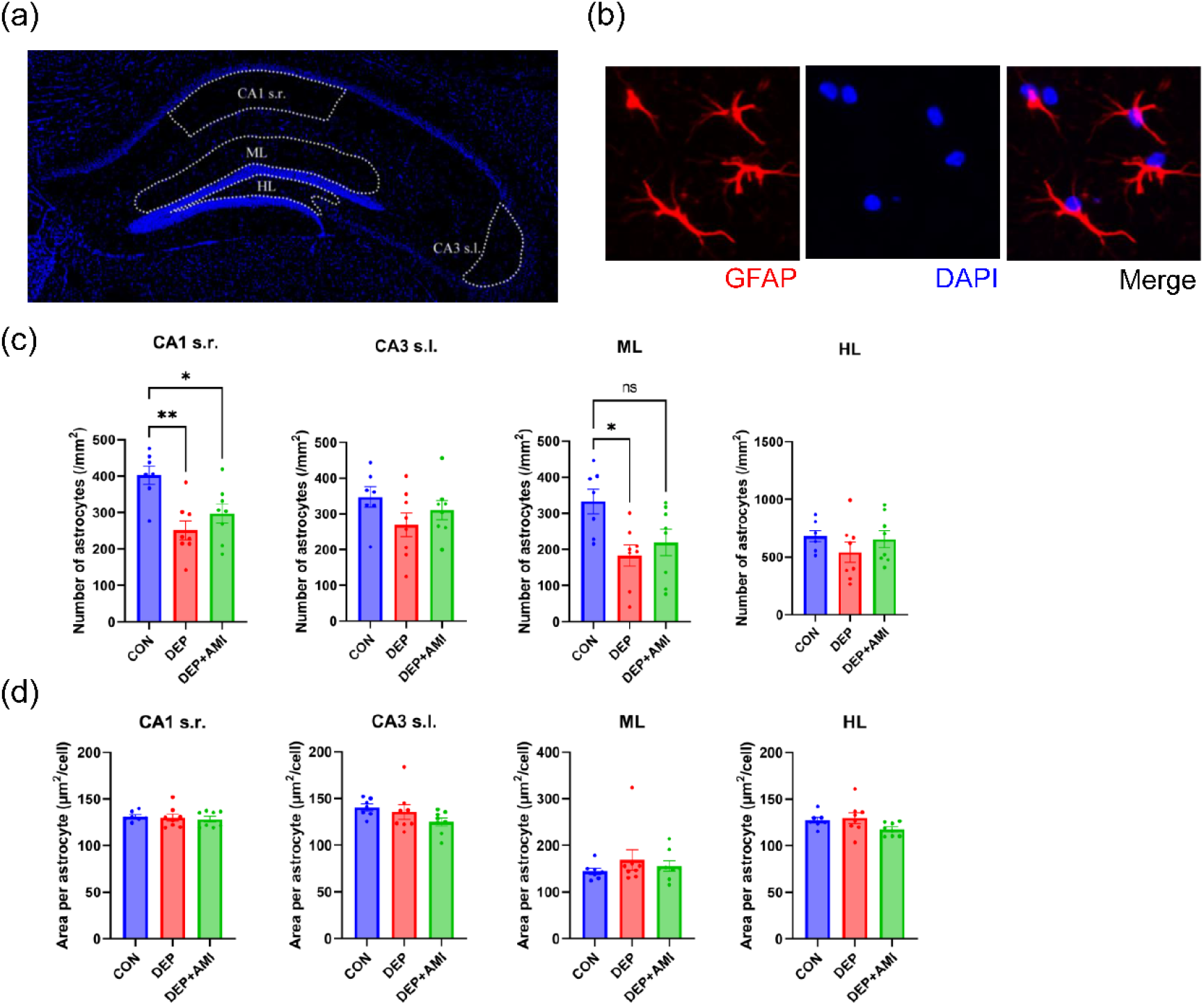
Decreased number of astrocytes in several subfields of the hippocampus in CORT-induced depression model mice. (a) Images of the entire hippocampus captured using a 40× objective, showing DAPI-positive nuclei. The regions of interest used throughout this study are outlined using white dotted lines. (b) Co-localization of GFAP and DAPI in the mouse hippocampus. (c) Quantification of GFAP-positive cells per mm^2^. n = 7–8 mice/group. The numbers of astrocytes in the CA1 s.r. and ML in the DEP group were lower than those in the CON group (*p* = 0.002 and *p* = 0.014, respectively; Tukey’s post hoc test). There was a similar decrease in the CA1 s.r. in the DEP+AMI group compared to that in the CON group (*p* = 0.026, Tukey’s post hoc test), but the difference was not as significant as in the DEP group. (d) Quantification of GFAP-positive area per cell. n = 7–8 mice/group. There were no significant differences between the groups in any region. CA1 s.r.: stratum radiatum region of the CA1, CA3 s.l.: stratum lucidum region of the CA3, HL: hilus, ML: molecular layer. ns, not significant; *, *p* < 0.05; **, *p* < 0.01.

### Isolation of hippocampal astrocytes by enzymatic dispersion and fluorescence-activated cell sorting (FACS)

At the end of the dosing regimen, the mice were euthanized by cervical dislocation. The hippocampus was dissected and a pair of hippocampi was used for each sample. Enzymatic cell dissociation and debris and red blood cell removal were performed using the Adult Brain Dissociation Kit (130-107-677; Miltenyi Biotec, Bergisch Gladbach, Germany) according to the manufacturer’s instructions. The dissociated single-cell suspension was resuspended and incubated in FcR Blocking Reagent (130–097-678; Miltenyi Biotec) at 4 °C for 10 min, and then co-stained for 15 min at 4 °C with PE-conjugated anti-astrocyte cell surface antigen-2 (ACSA-2) antibody (130-116-244; Miltenyi Biotec) and Alexa Fluor 647-conjugated anti-O1 antibody (FAB1327R; R&D Systems, Minneapolis, USA). To avoid sorting dead cells, 4′,6-diamidino-2-phenylindole (DAPI; 340-07971; Dojindo, Kumamoto, Japan) was also added immediately before sorting. The labeled cells were sorted using a SH800S Cell Sorter (Sony, Tokyo, Japan). One hundred ACSA-2 positive and O1 negative, ACSA-2 negative and O1 positive, and ACSA-2 negative and O1 negative cells each were collected in 1 µL aliquots of cell lysis buffer. Analyses were performed using the Cell Sorter software (v2.1.5) pre-loaded in the instrument; FlowJo software (v7.6.5; FlowJo LLC, OR, USA) was used for advanced data analysis.

### Reverse transcription with random displacement amplification (RT-RamDA)

The ACSA-2 positive and O1 negative, ACSA-2 negative and O1 positive, and ACSA-2 negative and O1 negative cells in cell lysis buffer were sorted directly into PCR tube strips. The cell lysis buffer was prepared using the QuantAccuracy RT-RamDA cDNA Synthesis Kit (RMQ-101; Toyobo, Osaka, Japan) according to the manufacturer’s protocol. The lysed cells were then used for cDNA synthesis using the same kit. Using RT-RamDA, both poly(A) and non-poly(A) RNAs, including nascent RNAs, histone mRNAs, long non-coding RNAs (lncRNAs), circular RNAs, and enhancer RNAs, can be detected with high sensitivity, while gDNA is removed and cDNA synthesis from rRNA is also prevented^18^. PCR was performed using the KOD FX polymerase (KFX-101; Toyobo); the primer sequences used are provided in Supplementary Table 1. Samples were analyzed after electrophoresis on a 3% agarose gel stained using MIDORI Green Xtra (MG10, NIPPON Genetics, Tokyo, Japan). For the positive control, RNA was extracted from the whole hippocampus using the AllPrep DNA/RNA Mini Kit (80204, QIAGEN), and RT-RamDA was performed using 100 cells amount of RNA.

### RNA-seq and sequence data analysis

Construction of a cDNA library using the RT-RamDA method^18^ was outsourced to the International Research Center for Medical Sciences, Kumamoto University. Sequencing of the cDNA library derived from the isolated hippocampal astrocytes was performed using the NextSeq 500 system (Illumina, San Diego, USA). Raw data from single-end sequences were cleaned using Trimmomatic (version 0.39) (LEADING:20, TRAILING:20, MINLEN:20, CROP:75)^19^ and checked for quality using FastQC. Filtered sequences were aligned to mouse reference sequences (GRCm38 primary assembly annotation) using the ultrafast RNA-seq aligner STAR (v2.7.9a).^20^ The resulting BAM files were input into RSEM (v1.3.3), which outputs the counts and normalized expression values (transcripts per million; TPM) of the sequencing reads for each gene and sample. The summed TPM values for each group and the top 75% of the genes with values greater than zero were included in the analysis. First, overlaps between the differential expression of two independent RamDA-seq comparisons were analyzed using the “stratified” rank-rank hypergeometric overlap (RRHO) method.^21, 22^ Then, to identify differentially expressed genes (DEGs), we used the TCC-GUI program^23^ with default settings^23, 24^; however, DESeq2 was used for normalization and DEG identification. Genes with fold change >= 2 and adjusted *p*-value <0.05 were selected as DEGs and submitted to the Ingenuity Pathway Analysis (IPA) software (QIAGEN, CA, USA). The IPA graphical summary gives an overview of the main biological findings of the analysis.

### Quantitative real-time reverse-transcription polymerase chain reaction (RT-PCR)

Total hippocampal RNA was extracted using the AllPrep DNA/RNA Mini Kit (80204, QIAGEN). RNA concentrations were measured using a spectrophotometer (NanoDrop Lite; Thermo Fisher Scientific). After reverse transcription into cDNA using the ReverTra Ace® qPCR RT Master Mix (FSQ-201; Toyobo), real-time PCR was performed using the StepOnePlus Real-Time PCR System (Applied Biosystems, CA, USA) and the THUNDERBIRD® SYBR® qPCR Mix (QPS-201, Toyobo) using appropriate primers (Supplementary Table 2). Samples were incubated at 95 °C for 30 s for initial denaturation, followed by 40 cycles of denaturation at 95 °C for 5 s and annealing and extension at 60 °C for 10 s. Relative changes in gene expression were calculated according to the comparative threshold cycle (CT) method and normalized to that of GAPDH as the internal control.

### Statistical analyses

All data are presented as mean ± SEM values, with statistical significance defined as *p* <0.05. Statistical analyses were performed using GraphPad Prism 9 (GraphPad Software, CA, USA). Results of experiments involving two groups were analyzed using the unpaired *t*-test. One-way analysis of variance (ANOVA) was used to compare the differences in means between groups in the univariate analysis. Two-way ANOVA was used in the multivariable analysis to model the relationship between the results of behavioral experiments, and CORT and AMI (independent variables). Interaction was tested in the model.

## Results

### Establishment of the mouse model of CORT-induced depression and assessment of antidepressant effects

We first confirmed the establishment of the mouse model of CORT-induced depression and evaluated the antidepressant effects of AMI using the SPT and NSFT (Fig. 1a). Changes in sucrose preference were examined (Fig. 1b), and it was found that the DEP group had significantly reduced sucrose preference (*p*<0.001) at 4 weeks (Fig. 1c). This reduction was recovered in the DEP+AMI group at 6 weeks (*p*=0.039) (Fig. 1d). In the NSFT, the DEP group had a significantly prolonged latency to feed (*p*<0.001), which was reduced in the DEP+AMI group (DEP☓AMI: *p*<0.001) (Fig. 1e, f). AMI treatment decreased travel distance in the OFT, but no interaction effects were observed between CORT and AMI administration (Supplementary Fig. 1). Blood CORT levels were significantly elevated in the CORT group (*p*<0.001) and were not altered by AMI treatment (*p*=0.275) (Fig. 1g); no interaction effects were observed (*p*=0.789). These results indicated that depression-like behaviors related to core clinical symptoms were induced in the DEP group, and that AMI treatment improved these behaviors independently of blood CORT levels and physical activity.

### Decreased number of astrocytes in several subfields of the hippocampalus subfields in CORT-induced depression model mice

We next examined how the number and size of hippocampal astrocytes were affected in the experimental mice. The levels of GFAP, a typical marker of astrocytes, in the entire hippocampus did not differ significantly among the CON, DEP, and DEP+AMI groups (Supplementary Fig. 2). However, astrocytic changes are not uniform in patients with MDD and vary according to the hippocampal subfield^12, 25^. Therefore, we evaluated the number and size of astrocytes in different hippocampal subfields using immunostaining for GFAP. Four synapse-rich hippocampal subfields that do not contain the main excitatory hippocampal neurons^26^ were selected, and the number and size of GFAP-positive astrocytes were examined (Fig. 2a, b). Compared to those in the CON group, the DEP group showed a significant decrease in astrocyte number in the stratum radiatum of the CA1 (CA1 s.r.) (*p*=0.002) and the molecular layer (ML) (*p*=0.014), but no significant changes in the stratum lucidum of the CA3 (CA3 s.l.) and the hilus (Fig. 2c). The DEP+AMI group showed a significant reduction in astrocyte number in the CA1 s.r. compared to that in the CON group, but to a lesser extent than the DEP group (*p*=0.026). There was no significant difference in the ML between the DEP+AMI and CON groups (*p*=0.069). Moreover, there were no significant differences in GFAP-positive area per astrocyte among the three groups in any of the four regions (Fig. 2d). Thus, the number of astrocytes was significantly decreased in the CA1 s.r. and ML in the DEP group, but there was no significant difference and a less significant difference, respectively, in the DEP+AMI group.

### Successful isolation of hippocampal astrocytes by enzymatic dispersion and FACS

Hippocampal astrocytes were isolated by enzymatic dispersion and FACS based on labeling with an anti-ACSA-2 antibody^27^ (Fig. 3a). As reported previously, since mature oligodendrocytes have low levels of Atp1b2, an ACSA-2 epitope, it was useful to distinguish oligodendrocytes using an anti-O1 antibody^28^, so we adopted that approach as well. Cells were first separated from debris based on the forward scatter area (FSC-A) and side scatter area (SSC-A) thresholds; then, double-labeled cells were removed by gating for FSC-A and forward scatter height (FSC-H). DAPI was used for excluding dead cells. ACSA-2-PE and O1-APC detection allowed the differentiation of four populations of live cells: ACSA-2-negative/O1-negative, ACSA-2-low positive/O1-positive, ACSA-2-low positive/O1-negative, and ACSA-2-positive/O1-negative (Fig. 3b). The latter three cell populations were isolated, and phenotypic purity was evaluated using RT-PCR. For some cell fractions, the number of cells collected per mouse was so low that normal RNA extraction was not possible. Therefore, RT-PCR was performed using the RT-RamDA method, which can produce cDNA from a small number of cells. ACSA-2-positive/O1-negative cells expressed astrocyte markers (Atp1b2 and Aqp4), but no other cell markers; ACSA-2 low positive/O1-positive cells specifically expressed the oligodendrocyte marker Mog; and ACSA-2 low positive/O1-negative cells specifically expressed the microglial marker Cx3cr1 (Fig. 3c). Therefore, we confirmed that astrocytes could be specifically isolated by isolating ACSA-2-positive/O1-negative cells.

**Figure 3.**
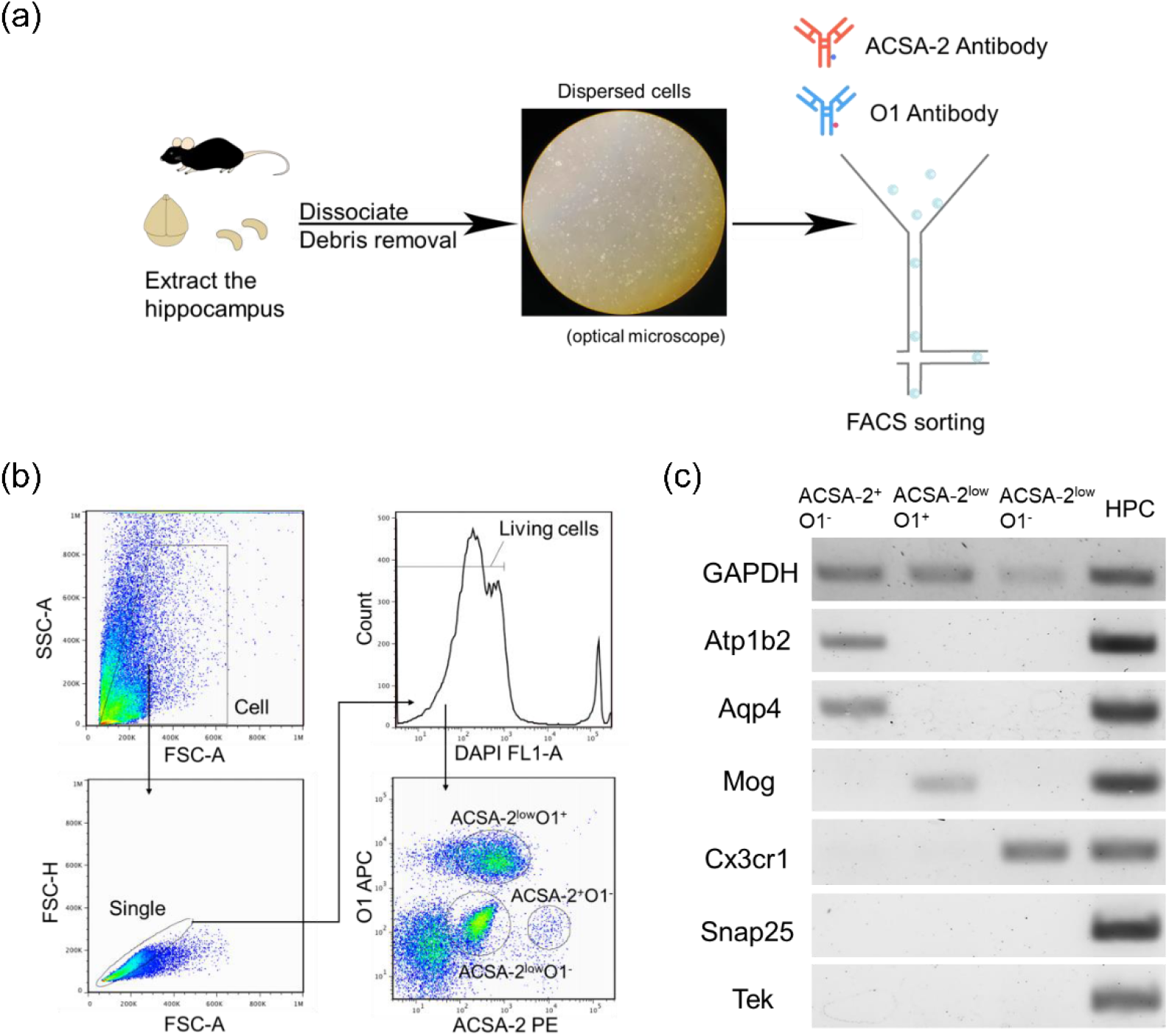
Successful isolation of hippocampal astrocytes by enzymatic dispersion and FACS. (a) Overview of the experimental manipulations to disperse and isolate hippocampal astrocytes. The image shows the cell lysate after dispersion under an optical microscope. (b) Gating strategy used for flow cytometry. Cells were initially differentiated from debris using FSC-A and SSC-A thresholds, with doublets subsequently excluded via FSC-A and FSC-H gating. Thereafter, DAPI was used for dead-live exclusion. This was followed by ACSA-2-PE and O1-APC detection, enabling the segregation of live cells into four distinct populations. Three of these regions were used for subsequent analysis. (c) The separation of cells was confirmed using RT-RamDA and PCR. For the positive control, RNA was extracted from the whole hippocampus and RT-RamDA was performed. Atp1b2 and Aqp4 are known markers of astrocytes, Mog for oligodendrocytes, Cx3cr1 for microglia, Snap25 for neurons, and Tek for vascular endothelial cells. The expected bands for astrocytes, oligodendrocytes, and microglia could be detected.

### AMI treatment induced opposite gene expression patterns in CON and DEP group hippocampal astrocytes

Using mice that exhibited the representative behaviors shown in Fig. 1, hippocampal astrocytes were isolated and RNA-seq was performed. RNA-seq data was initially evaluated using RRHO analysis to examine the relationships between the gene expression patterns altered by CORT and AMI treatments. RRHO analysis is a threshold-free approach that ranks all transcripts by *p*-value and direction of effect size, allowing the evaluation of the genome-wide overlap between two datasets (Fig. 4a).^21, 22^ The results showed that there was discordant overlap between the transcripts altered in the DEP group relative to the CON group and those altered in the DEP+AMI group relative to the DEP group (Fig. 4b), suggesting that the altered gene expression pattern in the DEP group may have been restored in the DEP+AMI group. Conversely, there was a harmonious overlap between the transcripts altered in the DEP group relative to the CON group and those altered in the CON+AMI group relative to the CON group (Fig. 4c). These results suggest that even if gene expression analysis is performed by administering antidepressants such as AMI to control mice, accurate analysis may be difficult because AMI treatment may result in different gene expression patterns depending on the pathology of the mice, as shown in the present results, thus reiterating the importance of using DEP mice.

**Figure 4.**
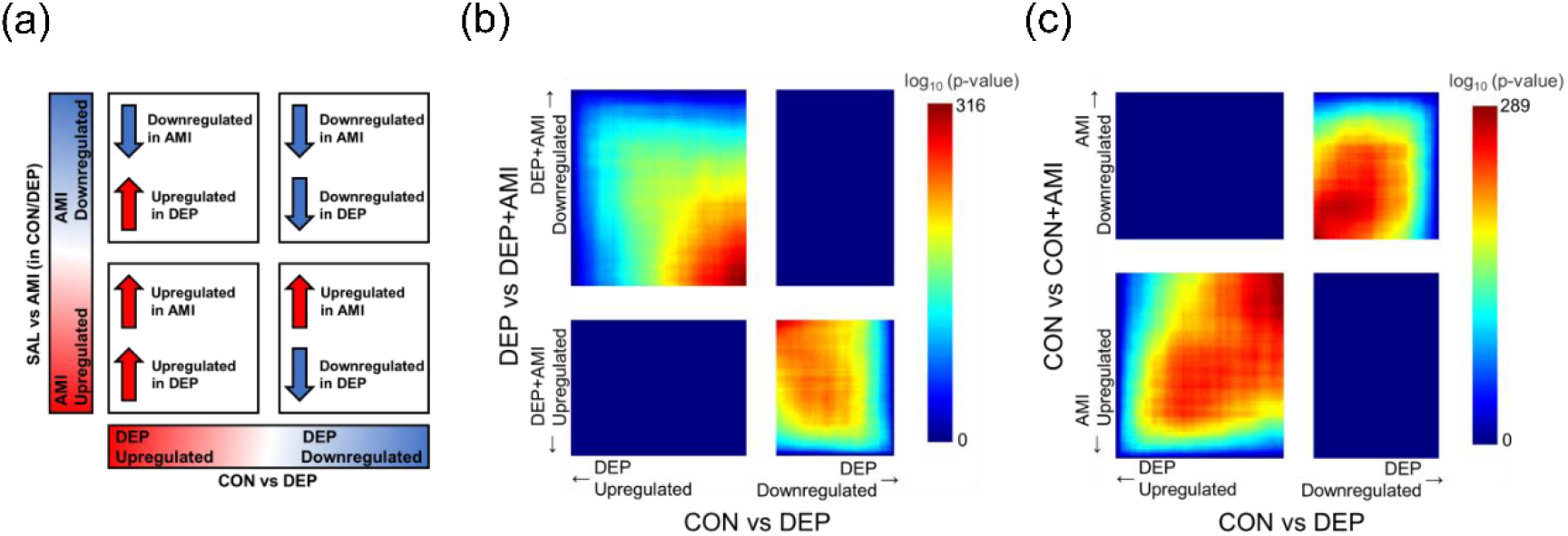
AMI treatment-induced opposite gene expression patterns of hippocampal astrocytes between the CON and DEP groups. (a) The stratified RRHO method involves a threshold-free approach that ranks genes by their *p*-value and effect size direction. Pixels represent the overlap between the transcriptome of each comparison as noted, with the significance of overlap [−log 10 (*p* - value) of a hypergeometric test] color coded. By using the RRHO test, it was possible to comprehensively assess the overlap by comparing the genes that varied between the two groups in each of the two studies. (b) Comparison of CON group versus DEP group with DEP group versus DEP+AMI group (max −log 10 (*p* -value) = 316). The RRHO results showed 18,808 genes across the four quadrants—upregulated in CON group (downregulated in DEP group) and downregulated in DEP group (upregulated in DEP+AMI group): 4249 genes; downregulated in CON group (upregulated in DEP group) and upregulated in the DEP group (downregulated in DEP+AMI group): 7687 genes. The results also showed that there was a discordant overlap between the transcripts altered in the DEP group relative to the CON group and those altered in the DEP+AMI group relative to the DEP group. (c) Comparison of CON group versus DEP group with CON group versus CON+AMI group (max −log 10 (*p* -value) = 289). The RRHO results showed 18,938 genes across the four quadrants—upregulated in CON group (downregulated in DEP) and upregulated in CON group (downregulated in CON+AMI group): 4750 genes; downregulated in CON group (upregulated in DEP group) and downregulated in CON group (upregulated in CON+AMI group): 7038 genes. The results showed that there was a harmonious overlap between the transcripts altered in the DEP group relative to the CON group and those altered in the CON+AMI group relative to the CON group.

### Identification of transcription factor 7 like 2 (TCF7L2) as a possible key regulator from among the oppositely regulated DEGs in the comparison between CON vs DEP and DEP vs DEP+AMI groups

As RRHO analysis suggested that variable genes in the DEP group are important, to identify genes involved in both pathogenesis and treatment, we searched for genes with variations in opposite directions among the DEGs between the CON vs. DEP and DEP vs. DEP+AMI groups. There were 553 DEGs between the CON and DEP groups (Fig. 5a), and 284 between the DEP and DEP+AMI groups (Fig. 5b). To evaluate the upstream regulatory genes of each DEG, we extracted the upstream regulators using the QIAGEN IPA software. The top ten regulators in order of absolute value of the activation z-score are shown in Fig. 5c and 5d. In comparing the CON and DEP groups, TCF7L2 was predicted as the most significant upstream regulator of reduced activity (z-score= −3.153) and had the lowest predicted probability of this outcome (-log_10_(*p*-value) = 8.883) (Fig. 5c). Fig. 5e provides further details regarding the TCF7L2-regulated DEGs depicted in Fig. 5c. Surprisingly, in the comparison between the DEP and DEP+AMI groups, only TCF7L2 was predicted to be active as an upstream regulator in the DEP+AMI group (z-score= 2.464) and also had the lowest predicted probability of this outcome (-log_10_(*p*-value) = 7.099) (Fig. 5d). Fig. 5f provides further details regarding the TCF7L2-regulated DEGs depicted in Fig. 5d. Since TCF7L2 was found to be the only upstream regulator whose activity was significantly decreased by DEP and recovered by AMI, we decided to examine the actual changes in TCF7L2 gene expression using qPCR and cDNA obtained from isolated astrocytes; however, due to low expression levels, it was difficult to achieve proper amplification (data not shown). TCF7L2 is reported to be most abundantly expressed in mouse hippocampal astrocytes.^29^ Therefore, we decided to use bulk hippocampal tissue for quantitative RT-PCR evaluation in this study. The results showed that TCF7L2 gene expression was significantly lower in the DEP group than in the CON group (*p* = 0.049) (Fig. 5g). In the DEP+AMI group, no significant reduction was observed compared to that in the CON group (*p* = 0.626).

**Figure 5.**
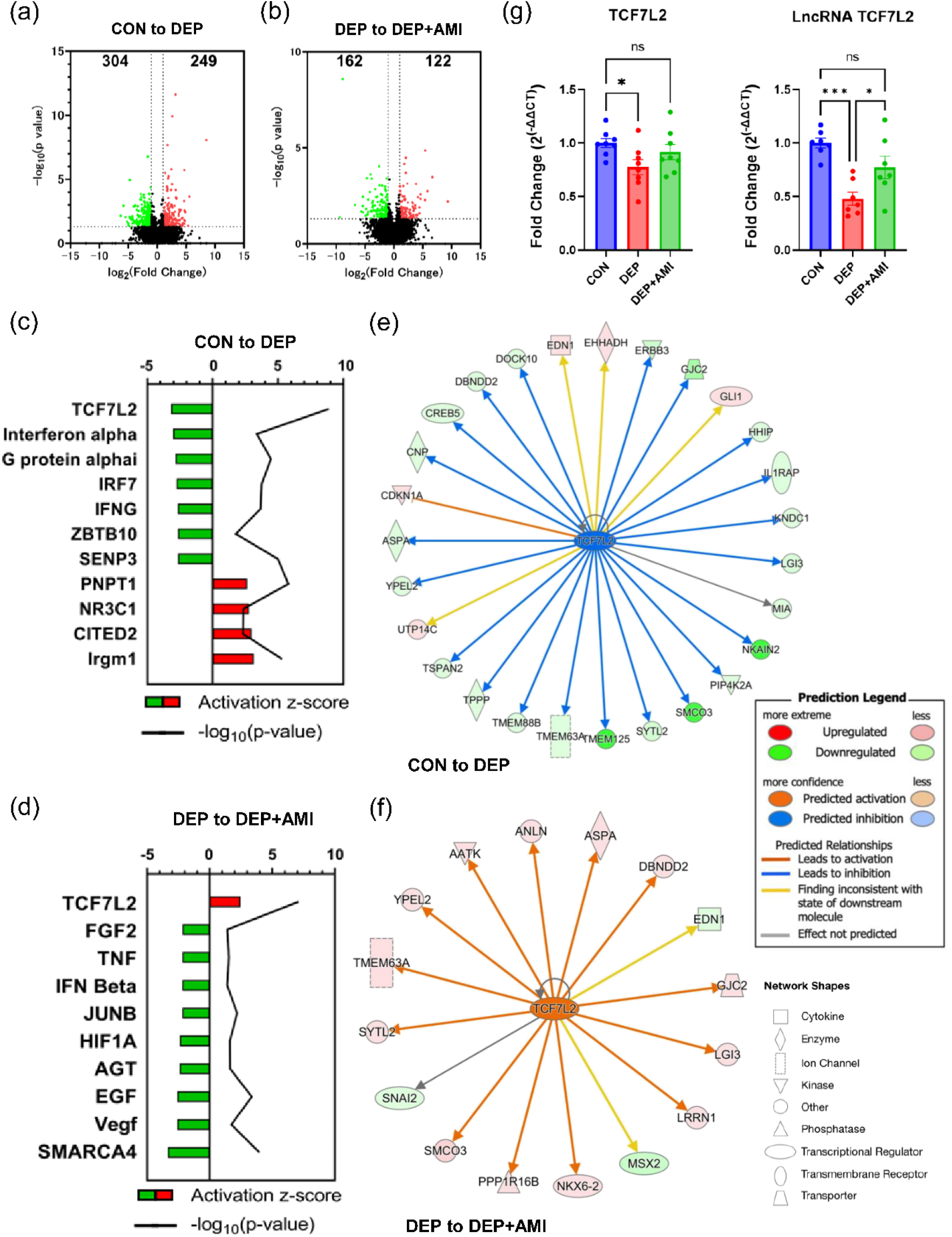
Identification of TCF7L2 as a key regulator among the oppositely regulating DEGs in the comparison between the CON vs DEP and DEP vs DEP+AMI groups. (a-b) Volcano plot displaying gene-wise log (fold change) on the x-axis against -log10(*p*-value) on the y-axis. The values shown in the figure are the number of DEGs (*p* < .05; |log2 (fold change)| > 1) for each. (a) Red dots indicate DEGs with increased gene expression in the DEP group (249 genes), and green dots indicate DEGs with decreased gene expression in the DEP group (304 genes). (b) Red dots indicate DEGs with increased gene expression in the DEP+AMI group (122 genes), and green dots indicate DEGs with decreased gene expression in the DEP+AMI group (162 genes). (c-f) Upstream analysis used the number of DEGs to predict upstream regulators of the biomarker genes. The Z-score was used to evaluate changes between each group. The top ten tied |Z-score| in each analysis are shown in the figure c, d. The line represents the *p*-value. (c) Using the 553 DEGs found between the CON and DEP groups, upstream analysis was performed. The DEP group predicted that TCF7L2 would be the most significant upstream regulator of reduced activity (z-score=-3.153) and had the lowest predicted probability of this outcome (-log10(*p*-value) = 8.883) (d) Upstream analysis was performed using the 284 DEGs found between DEP and DEP+AMI groups. The results showed that only TCF7L2 was predicted to be active as an upstream regulator in the DEP+AMI group (z-score= 2.464), and this group also had the lowest predicted probability of this outcome (-log10(*p*-value) = 7.099) (e) TCF7L2 and its downstream genes that were speculated to have altered activity between CON and DEP groups in this IPA analysis are shown. For example, the downregulation of TCF7L2 led to the inhibition (indicated by the blue arrow line) of the SMCO3 expression and the downregulation of SMCO3 is indicated by the green color. Other indicators are explained in the Prediction Legend section. (f) Comparing the DEP and DEP+AMI groups, this figure shows TCF7L2 and its downstream genes whose activity was altered in the DEP+AMI group. In contrast to the reduced activity of TCF7L2 in the DEP group compared to the CON group, TCF7L2 is activated and activates many downstream genes in the DEP+AMI group compared to the DEP group. (g) The gene expression of TCF7L2 and lncRNA TCF7L2 was assessed using qPCR in hippocampal tissue. n = 7–8 mice/group. TCF7L2 expression was found to be significantly downregulated in DEP group (*p* = 0.049, Tukey’s post hoc test). Regarding lncRNA TCF7L2, its expression was significantly decreased in DEP group (*p* < 0.001, Tukey’s post hoc test), and further, it significantly increased in DEP+AMI group (*p* = 0.032, Tukey’s post hoc test). ns, not significant; *, *p* < 0.05; ***, *p* < 0.001.

**Figure 6.**
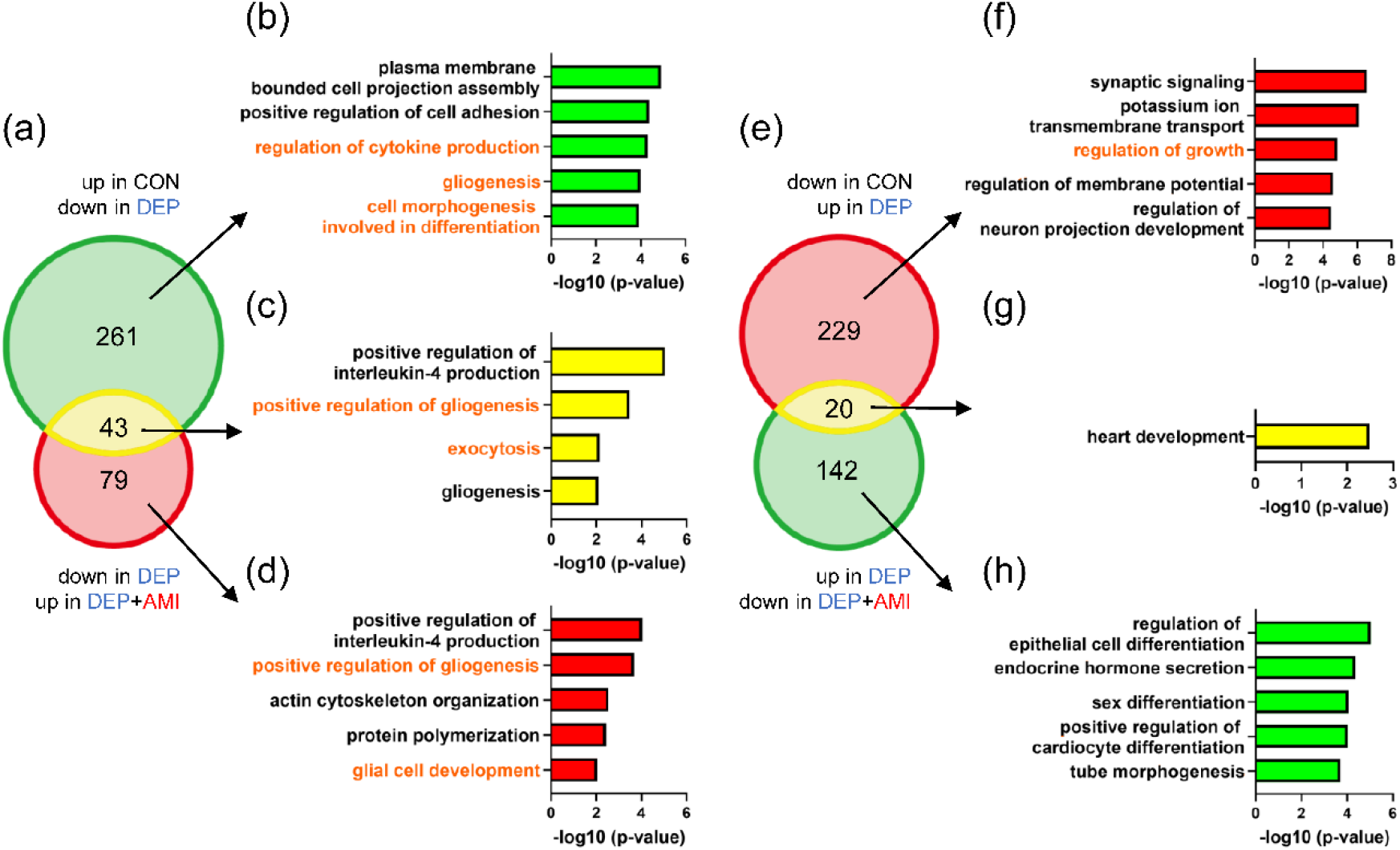
Biological processes of astrocytic TCF7L2 on in depression- and antidepressant-like behaviors. (a) The Venn diagram shows the overlap of the DEGs decreased in the DEP group between the CON and DEP groups and the DEGs increased in the DEP+AMI group between the DEP and DEP+AMI groups. (b-d) The top five biological processes in the GO analysis. The pathways in orange are those that contain genes downstream of TCF7L2 as pathway components (not included for genes indicated by IPA analysis as “ Finding inconsistent with state of downstream”). (e) The Venn diagram shows the overlap of the DEGs increased in the DEP group between the CON and DEP groups and the DEGs decreased in the DEP+AMI group between the DEP and DEP+AMI groups. (f-h) The top five biological processes in the GO analysis. The pathways in orange are those that contain genes downstream of TCF7L2 as pathway components (not included for genes indicated by IPA analysis as “Finding inconsistent with state of downstream”).

The TCF7L2 long non-coding RNA (lncRNA) is known to control the expression of its parent gene and is most abundant in astrocytes in the brain.^30^ Therefore, we also evaluated the expression of TCF7L2 lncRNA^30^. Expression of TCF7L2 lncRNA was also significantly lower in the DEP group than in the CON group (*p*<0.001). Additionally, it significantly recovered in the DEP+AMI group compared to that in the DEP group (*p*=0.032).

### Biological processes associated with astrocytic TCF7L2 in depression- and antidepressant-like behaviors

To evaluate how astrocytic TCF7L2 functions and influences depression-like behavior and antidepressant effects, Gene Ontology (GO) analysis was performed using Metascape (http://metascape.org).^31^ Fig. 5a shows the overlap between the DEGs downregulated in the DEP group and those upregulated in the DEP+AMI group. The top five biological processes in the GO analysis are shown in Fig. 5b-d. For each biological process, we checked whether the component genes included genes downstream of TCF7L2 shown in Fig. 4e and 4f (Supplementary Table 3). The biological processes shown in orange in Fig. 5b-d—regulation of cytokine production, gliogenesis, cell morphogenesis involved in differentiation, positive regulation of gliogenesis, exocytosis, and glial cell development*—*were identified as those that included downstream genes of TCF7L2. Conversely, Fig. 5e shows the overlap between DEGs upregulated in the DEP group and those downregulated in the DEP+AMI group. The top five biological processes in the GO analysis are shown in Fig. 5f-h; the biological process shown in orange—regulation of growth*—*was identified as containing genes downstream of TCF7L2. These GO analyses using RNA-seq data from isolated astrocytes suggest that the biological processes associated with TCF7L2 include proliferation, differentiation, and cytokine production.

## Discussion

In this study, we isolated hippocampal astrocytes and performed comprehensive gene expression analysis. Our two main findings were as follows. First, opposite gene expression patterns were observed in hippocampal astrocytes after AMI administration in depression model and control mice. This suggests that, even in the case of isolated astrocytes, it is important to use a mouse model of depression rather than normal mice to understand the molecular mechanisms of action of antidepressants. Second, TCF7L2 was the only gene differentially regulated in the context of both depressive and antidepressant-like behavioral changes in hippocampal astrocytes.

The importance of using mouse models of depression has long been noted. For example, chronic administration of antidepressants to normal mice is reported to induce anxiety-like behavior as a side-effect.^32^ Clinically, antidepressant administration to healthy individuals and patients with depression is reported to result in different changes in brain function.^33^ However, to the best of our knowledge, this is the first study to exhaustively examine the changes in gene expression in antidepressant-treated control and depression model mice. The results of our gene expression analyses support the validity of using model animals rather than normal animals, even when specific cell types are used, as in the present study.

TCF7L2 is an HMG box-containing transcription factor universally expressed in tissues throughout the body and plays an important role in the Wnt/β-catenin pathway, which is involved in cell proliferation and differentiation.^34–36^ TCF7L2 is also widely distributed in the central nervous system (CNS)^37, 38^, and is characterized by particularly high expression in astrocytes in humans and mice.^29, 30^ The Wnt/β-catenin pathway is well-known to regulate TCF7L2 activity^35^, but little is known about the mechanisms regulating TCF7L2 expression.

Recently, lncRNA-mediated regulation of mRNA expression has been reported as a novel regulatory mechanism of gene expression.^39^ Moreover, it has been reported that CORT reduces the expression of TCF7L2 lncRNA, which is abundant in the CNS, consequently leading to a decrease in the expression of the TCF7L2 gene.^30^ Our results also indicate that chronic CORT administration may induce reduced expression of TCF7L2 lncRNA in the hippocampus of DEP mice, potentially resulting in a reduction in TCF7L2 expression. However, since chronic AMI administration significantly increased TCF7L2 lncRNA expression in the hippocampus of DEP mice, it was assumed that TCF7L2 expression would also be significantly increased, but this was not the case. There can be two possible reasons for this. The first may be that using bulk tissue containing multiple cell types masked the changes in gene expression occurring in astrocytes, resulting in the failure to detect significant changes in TCF7L2 expression in astrocytes. The second reason may be that AMI restores depression-like behavior by activating TCF7L2 rather than restoring glucocorticoid-mediated decreased TCF7L2 expression. The fact that TCF7L2 was identified as a molecule that may be involved in the action mechanism of antidepressants by the IPA Upstream Regulator Analysis in this study, which estimated changes in activity rather than in expression, supports this possibility. To investigate these possible reasons, it will be necessary to develop a more efficient method of isolating astrocytes from the hippocampus in the future.

In this study, using a depression model and antidepressants, the physiological functions of TCF7L2 in hippocampal astrocytes were inferred to be proliferation, differentiation, and cytokine production using GO analysis. This suggests that the proliferation and differentiation of astrocytes and cytokine synthesis are impaired in the hippocampus of depression model mice. This would suggest that neuroplasticity, which has been reported to be associated with depression, is also impaired due to decreased astrocyte numbers, synapse formation failure, BBB disruption, and suppression of neurogenesis via cytokine synthesis. Moreover, as reported in previous basic and clinical studies, GFAP-positive astrocyte number is decreased in the hippocampal CA1 s.r. and ML of the dentate gyrus in depression model mice. This suggests that decreased TCF7L2 expression may be involved in suppressing the proliferative mechanisms of GFAP-positive astrocytes in these subregions.

Conversely, while AMI treatment seemed to promote the recovery of the number of GFAP-positive astrocytes in this study, the change was not statistically significant. This could be because our experimental system is not sensitive enough—the number of GFAP-positive cells does not necessarily reflect the number of astrocytes, since GFAP-positive cells represent only 15% of all astrocytes,^40^ and GFAP level reflects not only the number of astrocytes but also their activity.^41^ Another reason could be that daily intraperitoneal administration may have been highly stressful, resulting in smaller differences between the healthy and AMI-treated groups. Finally, glucocorticoids inhibit astrocyte proliferation, but antidepressants may promote differentiation of neural stem cells into astrocytes rather than restore astrocyte proliferation. Indeed, AMI increases the levels of various neurotrophic factors and induces the differentiation of neural progenitor cells.^2, 42^ Further investigation of these possibilities is expected to lead to a more detailed elucidation of the astrocyte-mediated action mechanism of antidepressants. Nevertheless, the present findings represent the results of a comprehensive analysis of the gene expression changes induced by CORT or AMI administration in hippocampal astrocytes, although it is still unclear whether these changes are directly related to the pathophysiology and treatment of MDD. It will be important to generate mice with site- and cell types-specific genetic modifications of TCF7L2 to investigate the pathophysiology of depression and the effects of antidepressants.

TCF7L2 is one of the major causative genes of type 2 diabetes mellitus (T2DM) and is associated with insulin metabolism.^35^ Recently, the high comorbidity of T2DM and mood disorders has attracted attention^40^, and two genetic polymorphisms of TCF7L2 have been reported to be associated with the risk of depression and T2DM.^41^ Furthermore, another genetic polymorphism of TCF7L2 has been shown to be associated with the risk of T2DM in bipolar disorder.^42^ Thus, decreased TCF7L2 function is likely to be associated with depression centrally, probably by causing abnormalities in astrocyte proliferation, differentiation, and cytokine production, as suggested in the present study, and peripherally, through decreased insulin secretion, which may contribute to T2DM via reduced peripheral insulin levels. In other words, common dysfunctions of TCF7L2 may be involved in the pathogenesis of both mood disorders and T2DM in different organs and through different mechanisms. In the same chronic CORT administration mouse model of depression used in this study, depression-like behavior as well as a higher insulin resistance index and increased insulin levels, that is, a precursor state to T2DM, has also been reported.^40^ Thus, persistent corticosterone overstimulation may lead to systemic TCF7L2 dysfunction, concomitant mood disorders, and T2DM; however, further studies are required to analyze these aspects in detail.

A possible limitation of this study is that only male mice were used. There is considerable evidence that females are more likely to have depression than males, although there is no adequate explanation for this sex-related difference.^43^ Thus, the involvement of TCF7L2 in female mice is of great concern and remains a subject for future studies. Another possible limitation is that AMI was the only antidepressant used in this study; therefore, the results cannot be generalized to other antidepressants.

In conclusion, we identified TCF7L2 as a gene associated with depression-like and antidepressive behaviors in response to AMI in hippocampal astrocytes, possibly via cellular physiological functions such as proliferation and cytokine production. These findings could provide valuable insights into the mechanisms underlying astrocyte-mediated antidepressant effects.

## Acknowledgments

We thank K. Iwamoto and Y. Nakachi (Kumamoto University) for helpful comments on this manuscript; S. Usuki (Kumamoto University) for technical support regarding RNA sequencing; and N. Maeda, K. Matsuura, and K. Okamura (Kumamoto University) for technical assistance with the experiments. We also thank Maanya and the other staff from Editage (www.editage.com) for carefully proofreading the manuscript. This work was supported by JSPS KAKENHI (Grant Number: 18H02756, 23H02839), Takeda Science Foundation, and the program of the Joint Usage/Research Center for Developmental Medicine, Inter-University Research Network for Trans-Omics Medicine, Institute of Molecular Embryology and Genetics, Kumamoto University.

## Supporting Information

### Behavioral tests

The sucrose preference test (SPT) was performed weekly. The novelty suppressed feeding test (NSFT) and open field test (OFT) were performed in week 7. Before the SPT, mice were given access to 1% aqueous sucrose solution for 72 h to allow them to adapt to the sucrose. After adaptation, mice were deprived of water for 24 h and then given access to two standard drinking bottles, one containing 1% sucrose and the other containing tap water, for 24 h. The positions of both bottles were switched at 0.5, 2, 4, and 8 h to account for any preference for a particular side. Sucrose and water intakes were measured at 0.5 and 24 h. Sucrose preference was calculated as follows: sucrose preference (%) = sucrose intake / (sucrose intake + water intake) × 100.

The NSFT was performed after the mice were food-restricted for 24 h. During the test (900 s), a mouse was placed in the same corner of an open-field apparatus (50 × 50 × 40 cm) in which a food pellet was placed centrally on a piece of white paper (9 × 9 cm). Latency was defined as the time from the placement of the mouse in the apparatus to that of its first bite of food.

The OFT was performed under bright interior lighting to evaluate the locomotor and exploratory behaviors of the mice. The floor of this open-field apparatus (50 × 50 × 40 cm) was divided into nine equal squares, and its color and texture were different from those of the apparatus used in the NSFT. The mouse was placed in a corner area and then the total distance moved was recorded for over 5 min. Each movement distance was calculated using SMART (version 3.0.06) based on the recorded video.

### Western blotting analysis

Whole hippocampus samples were homogenized in RIPA lysis buffer (50 mM Tris-HCl [pH 8.0], 150 mM NaCl, 1% Nonidet P-40, 0.5% deoxycholic acid, 0.1% sodium dodecyl sulfate [SDS], and 1 mM EDTA) containing a protease inhibitor cocktail (04080-24; Nacalai Tesque Inc., Kyoto, Japan) and phosphatase inhibitors (07575-51, Nacalai Tesque). The supernatant was collected by centrifugation at 10,000 rpm for 20 min at 4 °C, and total protein concentration was determined using a Pierce BCA Protein Assay Kit (Thermo Fisher Scientific). Equal amounts of protein (1 μg) for each sample were loaded onto a 12% SDS-polyacrylamide gel electrophoresis gel for electrophoresis. Polyvinylidene difluoride (PVDF) membranes (#1620177; Bio-Rad Laboratories, CA, USA) with the transferred proteins were blocked with 5% skim milk in Tris-buffered saline with Tween 20 (TBST) and then incubated first with antibodies against GFAP (1:1000; sc-58766; Santa Cruz Biotechnology, Dallas, USA) or Erk1/2 (1:2000; #4695; Cell Signaling Technology, Danvers, USA) as loading control overnight at 4 °C. The next day, the membranes were washed with TBST (5 × 5 min) and incubated with the corresponding horseradish peroxidase (HRP)-linked secondary antibodies for 1 h at room temperature. The blots were washed five times with TBST, and protein bands were detected using a chemiluminescence detection reagent (#170-5060; Bio-Rad Laboratories). The optical densities of the protein bands were estimated using Image Lab 6.1 (Bio-Rad Laboratories) and normalized to those of Erk1.

### Immunohistochemistry and imaging

Mice were sacrificed by transcardial perfusion with 20 ml of phosphate-buffered saline (PBS) containing heparin (10 U/ml; 17513-41; Nacalai Tesque) and verapamil (0.2 mg/ml; 222-00783; Wako, Osaka, Japan), followed by 20 ml of 4% PFA. The brains were post-fixed overnight in 4% PFA at 4 °C and subsequently cryo-protected overnight in 30% sucrose solution at 4 °C. Brains were embedded in OCT compound (45833; Sakura Finetek, Tokyo, Japan) and frozen at −80 °C. Frozen samples were sectioned (thickness, 10 μm) using a cryostat (CM1950; Leica, Wetzlar, Germany) and mounted onto glass slides. For immunofluorescence assays, the sections were permeabilized with 0.01% Triton X-100 in PBS (PBST) for 30 min. Antigen retrieval was performed by boiling the slides in 0.01 mol/L citrate buffer (pH 6.0) for 10 min. After washing twice with PBST, the sections were preincubated with 3% bovine serum albumin (BSA; 01281-26; Nacalai Tesque) in PBST at room temperature for 1 h. Slides were then incubated overnight at 4 °C with primary antibody (mouse anti-GFAP; SC-58766; Santa Cruz Biotechnology, Dallas, USA; 1:1000). Sections incubated with the antibody diluent without primary antibody were used as negative controls. After washing thrice with PBST, the slides were incubated with the appropriate secondary antibody (goat anti-mouse IgG1-Alexa 647; 1:1,000; 115-607-185; Jackson, PA, USA) for 1 h at room temperature. Nuclei were stained with 4′,6-diamidino-2-phenylindole (DAPI; 340-07971; Dojindo, Kumamoto, Japan), and the slides were then washed three times in PBST before imaging.

---

**Supplementary Figure 1.**
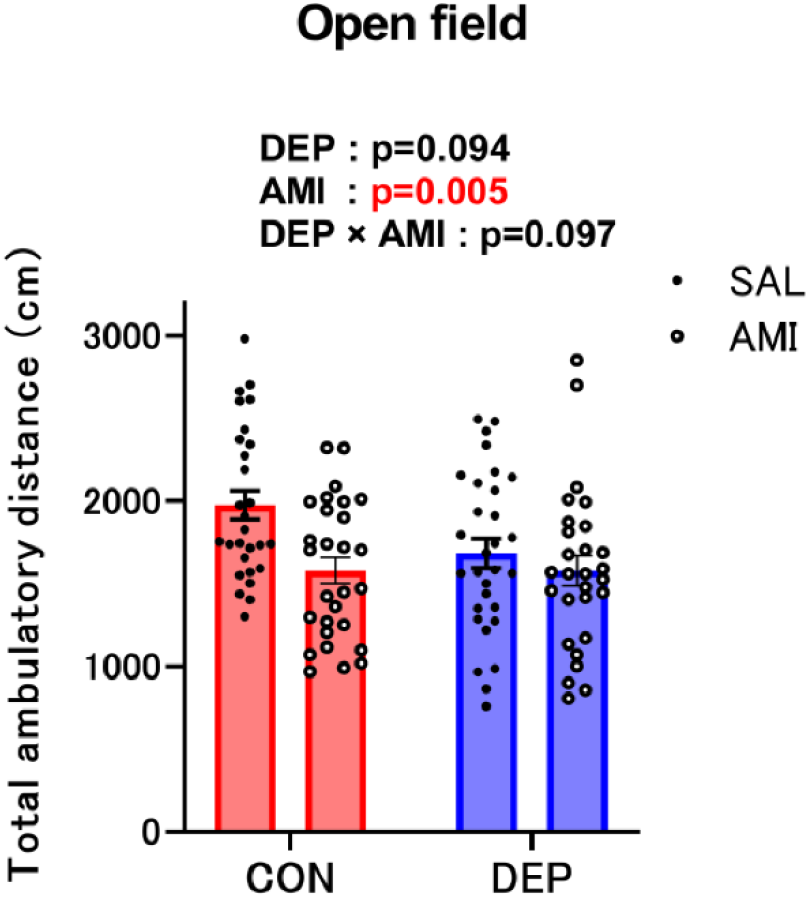
The result of the OFT conducted to assess locomotor activity. The OFT to confirm that CORT and AMI were not exerting a strong influence on the animals’ activity levels revealed that AMI treatment decreased travel distance (F (1, 110) = 8.348, *p =* 0.005, two-way ANOVA), but no interaction effects were observed between CORT and AMI administration (DEP☓AMI: F (1, 110) = 2.800, *p* = 0.097, two-way ANOVA).

**Supplementary Figure 2.**
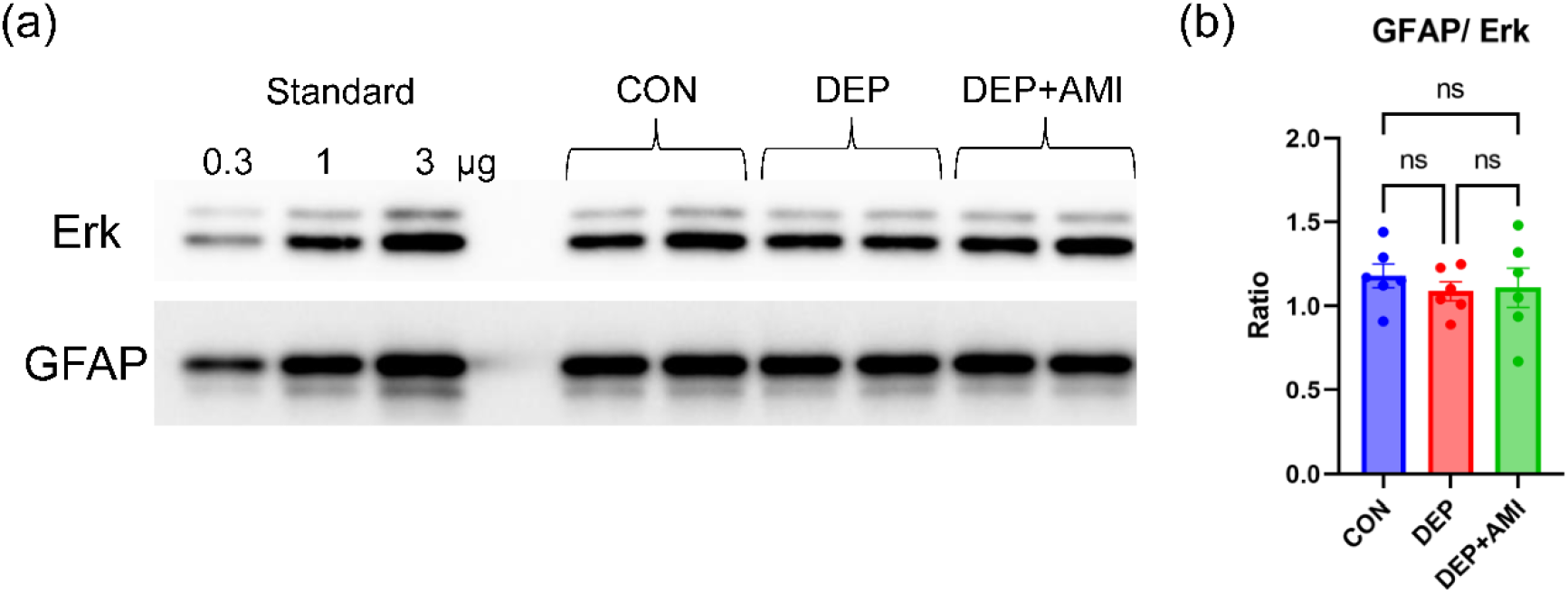
GFAP protein level in the entire hippocampus. (a) Western blot of GFAP protein from CON,DEP, and DEP+AMI groups. Erk was used as a loading control. (b) There did not differ significantly among the CON, DEP, and DEP+AMI groups (F (2, 15) = 0.319, *p* = 0.732, one-way ANOVA). ns, not significant.

**Supplementary Table 1.**
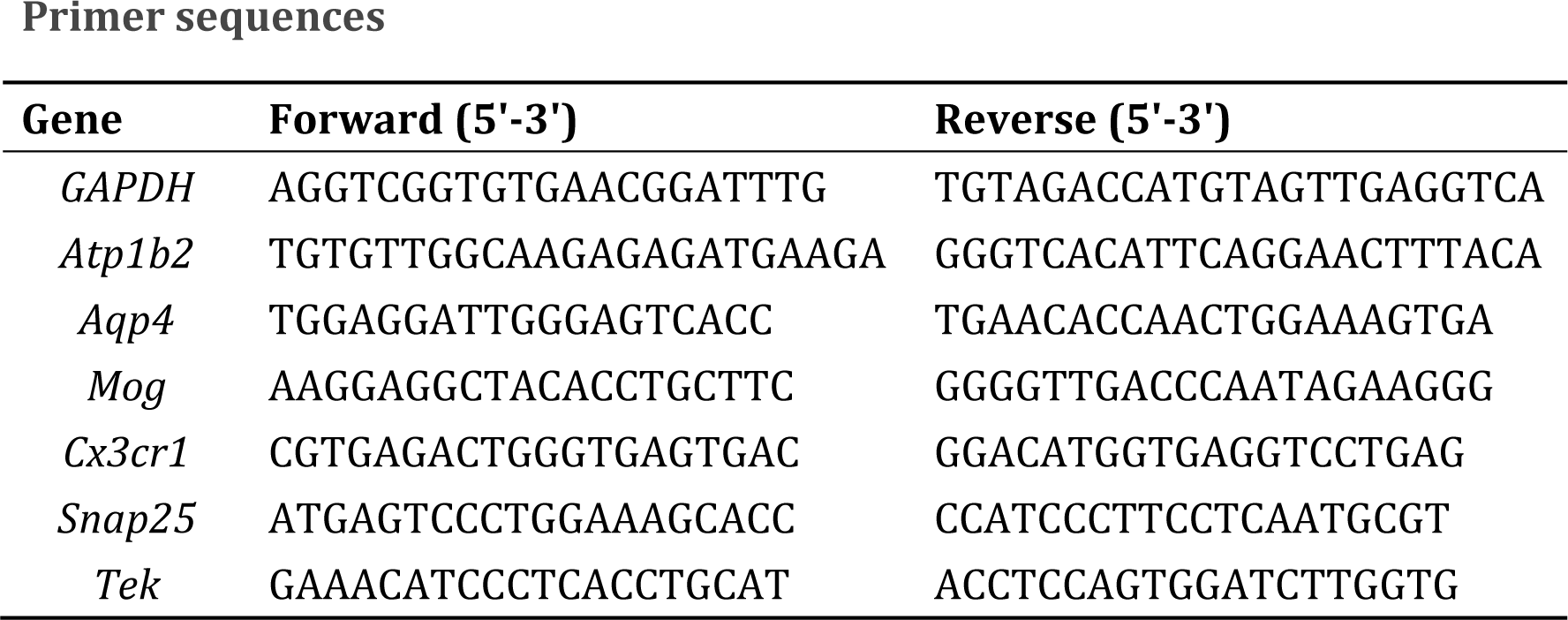

**Supplementary Table 2.**
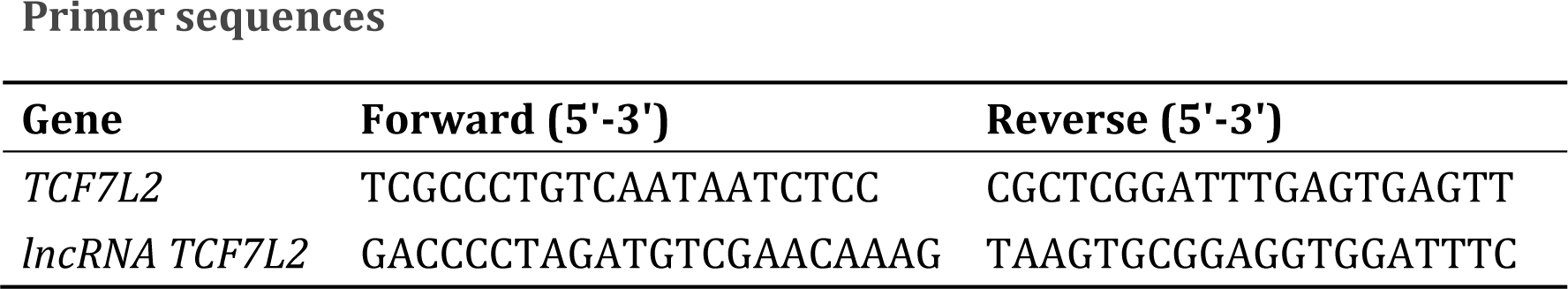

**Supplementary Table 3.**
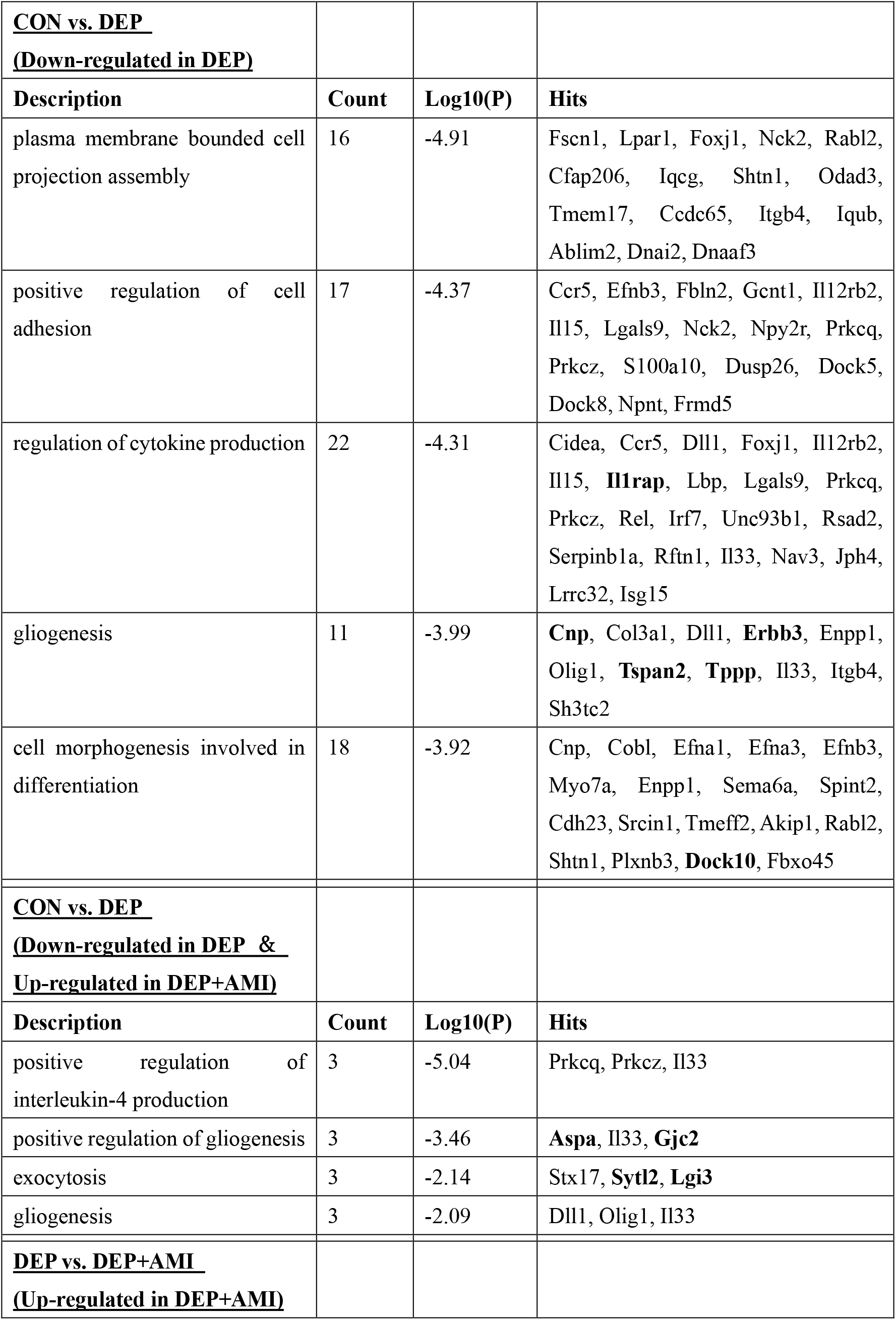
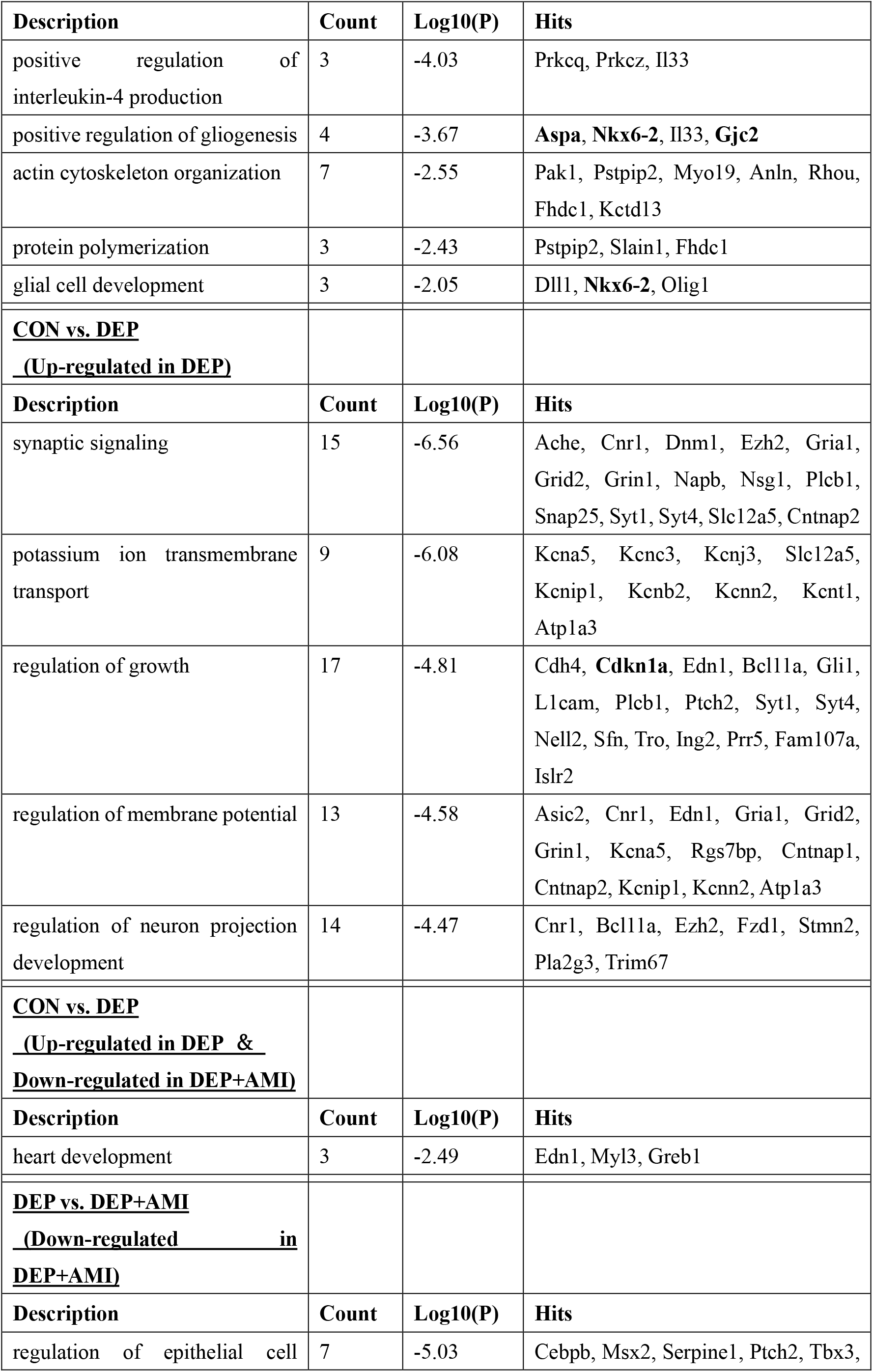
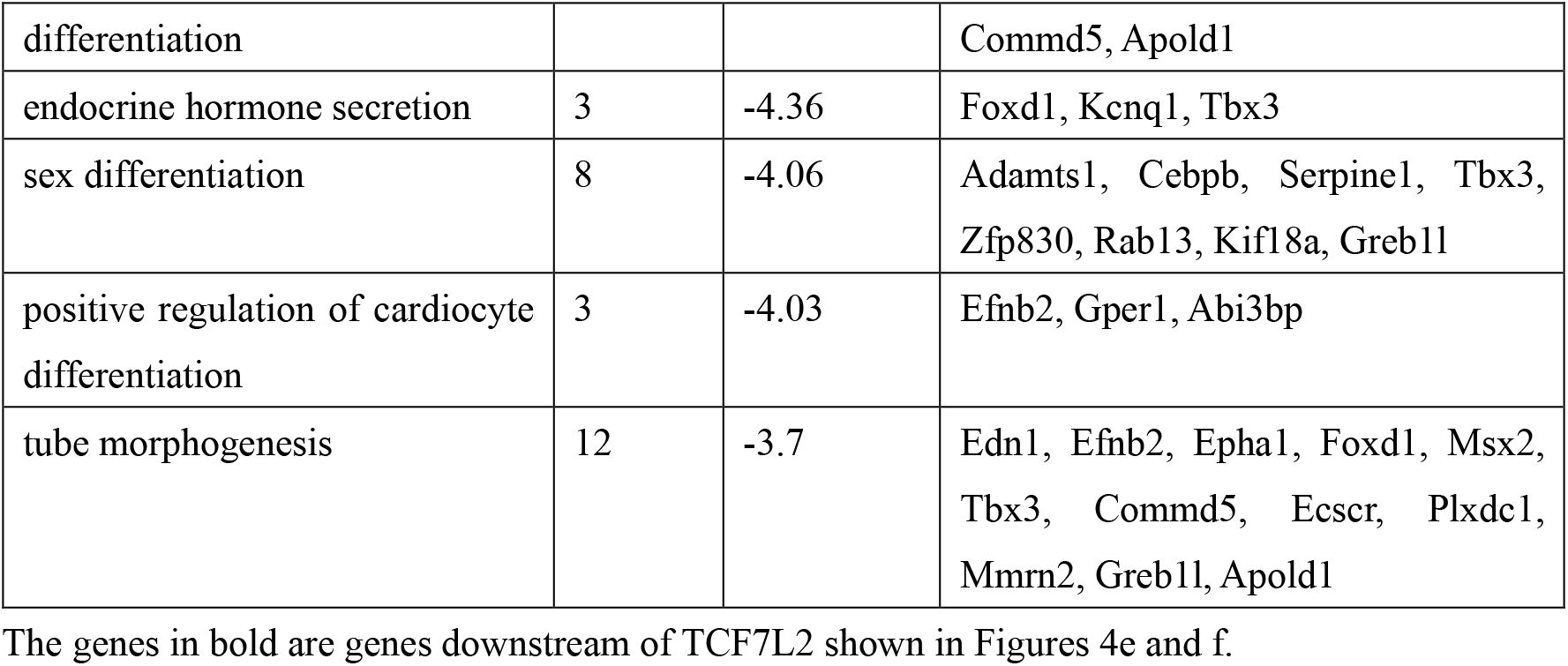

## Notes

**Disclosure Statement:** The authors declare no conflicts of interest relevant to this study.

### Competing Interest Statement

The authors have declared no competing interest.

